# Aquaporin-4 facilitator TGN-073 demonstrates novel analgesic activity

**DOI:** 10.1101/403311

**Authors:** Vincent J. Huber, Ingrid L. Kwee, Tsutomu Nakada

**Affiliations:** Center for Integrated Human Brain Science, Brain Research Institute, University of Niigata, Niigata, Japan.; Department of Neurology, University of California, Davis, California, USA.

## Abstract

During pre-clinical development, we tested the novel, internally developed AQP-4 facilitator TGN-073 for its effect in a rodent pain model. Therein, TGN-073 was found to exert a strong analgesic effect. Following a single 200 mg/kg (i.p.) administration of TGN-073, a virtually complete block in the acetic acid writhing test was observed. Subsequent *in vitro* tests demonstrated that TGN-073 had no binding affinity for the μ-opioid or NK-1 receptors. Accordingly, we suspect TGN-073 or other AQP-4 facilitators may be developed into potent non-opioid analgesic agents. Given the potential significance of this discovery, we feel it should be openly shared with the scientific community.

## Introduction

Aquaporin-4 (AQP-4) is an integral membrane water channel known to have a specific distribution in the central nervous system (CNS),^1^ which has led to considerable interest in its potential as a pharmacological target.^2^ Inhibitors of AQP-4 water transport were considered to be the most pharmacologically relevant,^3–4^ but there is now evidence suggesting potential roles for facilitators, compounds that increase AQP-4 water flux, in the treatment of neurodegenerative diseases. Our laboratory recently identified TGN-073 (3-benzyloxy-2-phenylsulfonamido-pyridine) as a pharmacologic AQP-4 facilitator that can be used to test the effects of increased AQP-4 function *in vivo*.^5^ However, reduced pain stimuli response was observed in AQP-4 null rodents,^6–7^ and an analgesic effect was reported for the AQP-4 inhibitor TGN-020 in a rodent neuropathic pain model.^8^ Consequently, we were concerned that AQP-4 facilitators, such as TGN-073, could promote a neurogenic pain syndrome, and felt it essential to test such compounds in an appropriate rodent pain model.

## Results and Discussion

Following approval by the Internal Review Board of the University of Niigata, we outsourced the testing of TGN-073 to Eurofins Pharmacology Discovery Services Taiwan in a blinded fashion. The results are shown in the accompanying figure. Single administration of TGN-073 sodium salt (200 mg/kg, i.p.) led to a virtually complete block in the writhing test. To our knowledge, few agents exhibit this level of analgesia in the standardized acetic acid writhing test other than opioids. Follow up *in vitro* tests, outsourced to Eurofins Panlabs Discovery Services, demonstrated that TGN-073 had no significant binding affinity for the μ-opioid and NT-1 receptors, 0 and 12%, respectively at 10 μM ligand concentration. Furthermore, interaction of TGN-073 with COX-2 was not suspected because of structural considerations, and was consequently not tested.

Accordingly, we suspect TGN-073 or other AQP-4 facilitators have significant potential to be developed into non-opioid analgesic agents. Given that we are not pain researchers and our laboratory is not equipped for such research, we strongly believe we should not pursue this line of investigation for academic purposes. Nevertheless, we felt that this discovery should not be buried, rather it should be openly shared with the scientific community, including those who can build on this result.

**Figure.**
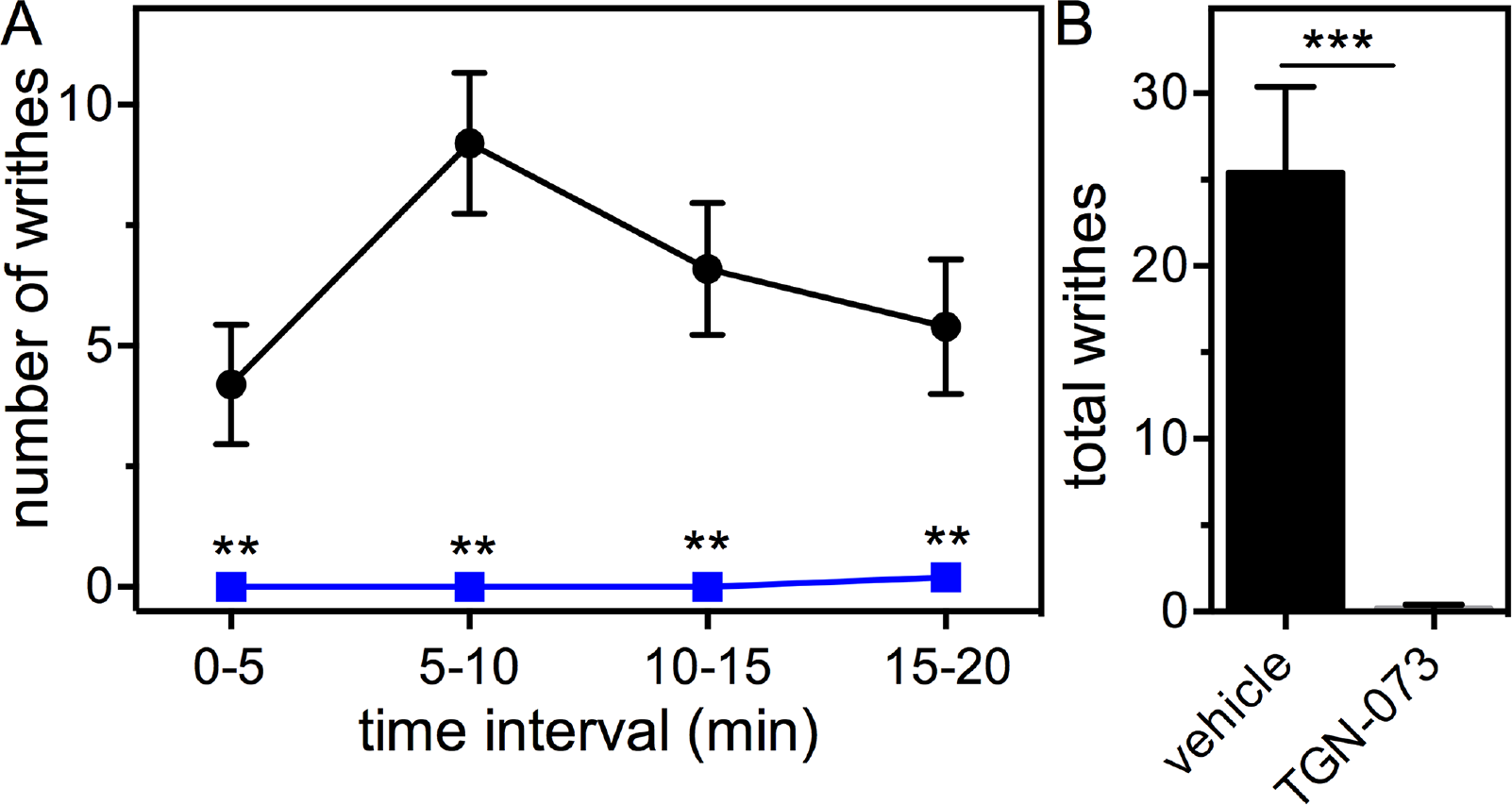
(A) TGN-073-Na (200mg/kg) or a normal saline vehicle (0.9% NaCl 5mL/kg) were administered intraperitoneally (i.p.) to groups of 5 ICR derived male mice (BW = 23 ± 3 g) 30 minutes before injection of acetic acid (0.5%, 20mL/kg, i.p.). The number of writhes were recorded in 5-minute intervals for 20 minutes total. Blue squares: TGN-073, Black circles: vehicle, *n* = 5, mean ± SE, ** p < 0.01 (Two-way ANOVA followed by Bonferroni correction). (B) Total writhes over the full 20 minutes. Mean ± SE. ***p <0.001 (Two tailed t-test).

## Materials and Methods

### Test compound

TGN-073 was obtained in bulk form from Key Organics, Ltd (Camelford, Cornwall, UK). The material obtained was found to be chemically identical to an in-house synthesized sample,^5^ and was consistent with having a chemical purity >95% according to ^1^H-NMR and UPLC-HR-MS analysis. The sodium salt for *in vivo* administration was prepared as described in Reference 5, and the resulting material was used as thusly obtained.

### In vivo pain model

Following approval by the Internal Review Board of the University of Niigata, TGN-073 was tested in the acetic acid writhing test (catalog number 503900) outsourced to Eurofins Pharmacology Discovery Services Taiwan, Ltd (Taipei, Taiwan) in a blinded fashion. Therein a single dose of TGN-073 sodium salt (200 mg/kg) in normal saline (5 mL/kg) or the same volume of vehicle was administered by intraperitoneal (i.p.) injection 30 minutes prior to an acetic acid challenge (0.5%, 20 mL/kg, i.p.) in *n* = 5 ICR derived mice per test and control groups, respectively. Following the acetic acid challenge, the number of writhes were counted in five-minute intervals for 20 minutes. Statistical significance was determined by Two-way ANOVA followed by Bonferroni correction.

### In vitro receptor binding

Assessment of TGN-073 binding with the μ-opioid and NK-1 receptors was outsourced to Eurofins Panlabs Discovery Services Taiwan (Taipei, Taiwan). TGN-073 (10 μM) was tested in radioligand binding assays for μ-opiate receptor (catalog number 260410), and Tachykinin NK-1 receptor (catalog number 255520) for *n* = 3 replicates.

## Acknowledgements

Funding was provided by the Ministry of Education, Culture, Sports, Science and Technology, Japan.

